# Cross-border investigations on the prevalence and transmission dynamics of *Cryptosporidium* species in dairy cattle farms in western mainland Europe

**DOI:** 10.1101/2021.10.18.464852

**Authors:** Pedro Pinto, Cláudia A. Ribeiro, Sumaiya Hoque, Ourida Hammouma, Hélène Leruste, Sébastien Detriche, Evi Canniere, Yvonne Daandels, Martine Dellevoet, Janine Roemen, Anne Barbier Bourgeois, Martin Kváč, Jérôme Follet, Anastasios D. Tsaousis

## Abstract

*Cryptosporidium* is comprised an apicomplexan parasitic protist, which infects a wide range of hosts, causing cryptosporidiosis. In cattle farms, the incidence of cryptosporidiosis results in high mortality in calves leading to considerable economic loss in the livestock industry. Infected animals may also act as a major reservoir of *Cryptosporidium* spp., in particular *C. parvum,* the most common cause of cryptosporidiosis in calves. This poses a significant risk to other farms via breeding centres, to trading of livestock and to human health. This study, funded by the Interreg-2-seas programme, is a part of a global project aimed at strategies to tackle cryptosporidiosis. To reach this target, it was essential to determine whether prevalence was dependent on the studied countries or if the issue was borderless. Indeed, *C. parvum* occurrence was assessed across dairy farms in certain regions of Belgium, France and the Netherlands. At the same time, the animal-to-animal transmission of the circulating *C. parvum* subtypes was studied. To accomplish this, 1084 faecal samples, corresponding to 57 dairy-farms from all three countries, were analysed. Well-established protocols amplifying the *18S* rDNA and *gp60* genes fragments, followed by DNA sequencing, were used for the detection and subtyping *C. parvum*; the DNA sequences obtained were further characterised using a combination of bioinformatics and phylogenetics methods. Our results show 25.7%, 24.9% and 20.8% prevalence of *Cryptosporidium* spp. in Belgium, France and the Netherlands respectively. Overall, 93% of the farms were *Cryptosporidium* positive. The *gp60* subtyping demonstrated a significant number of the *C. parvum* positives belonged to the IIa allelic family, which has been also detected in humans. Consequently, this study highlights how widespread is *C. parvum* in dairy farms and endorses cattle as a major carrier of zoonotic *C. parvum* subtypes, which subsequently pose a significant threat to human health.

## Introduction

*Cryptosporidium* is a genus of enteric apicomplexan parasites, responsible for causing cryptosporidiosis in a diverse range of vertebrate hosts, including livestock and humans [1, 2]. Cryptosporidiosis is one of the prominent causes of diarrheal illness in humans and animals worldwide, with children, newborn animals and immunocompromised individuals being especially vulnerable to the disease [3–5]. *Cryptosporidium* infections occur after ingestion of oocysts through the fecal-oral route, either directly after contact with infected animals or indirectly through contaminated material such as food, water, soil and fomites [6, 7]. Thus, an One-Health approach, whereby all these factors are considered, is essential to investigate the role and transmission dynamics of this parasite in both humans and other animals [7].

In livestock, cryptosporidiosis primarily manifests as a gastrointestinal disease, causing watery diarrhea, malnutrition, abdominal pain, dehydration and, in severe cases death [7, 8]. This disease is deemed globally endemic in cattle and it is particularly prevalent in neonatal and pre-weaned calves (< 6 weeks old), in which it is one of the most common causes of diarrheic illness [9, 10]. *Cryptosporidium* infection and consequent disease appear in both beef and dairy calves, with higher prevalence in intensive livestock management systems [11]. Clinical manifestations of cryptosporidiosis in calves lead to long term impacts on their well-being and consequently, the disease puts a considerable economic burden on the cattle industry [9, 10, 12]. Costs associated with the economic burden of the disease include: seeking veterinary expertise for diagnosis and medication, along with additional costs linked to animal rearing and supplemental nutrition to regain meat and milk yield. An additional cost of purchasing new animals has to be incurred in case of death [13]. In the United Kingdom, the extra cost to manage diarrheal diseases, including *Cryptosporidium*, has been estimated to an average of £32 per calf with an annual total spending of £11 million [13, 14]. More recently the economic burden of cryptosporidiosis in calves was highlighted, with projections reaching £100-200 per *Cryptosporidium*-infected calf. The long-term difference in growth of beef calves with and without cryptosporidiosis was also assessed [10, 15]. An average variation of 34 kg was observed between the two groups, with infected calves being considerably lighter than the uninfected, translating into a profit deficit of approximately £128 per animal according to market prices in 2018 [10]. Hence, studying the prevalence and transmission dynamics of this parasite in cattle farming is crucial.

*Cryptosporidium* spp. incidence in cattle is mostly attributed to four species: *C. parvum, C. bovis, C. andersoni* and *C. ryanae* [3]. Remarkably, an age-related distribution pattern has been noted, with *C. bovis* and *C. ryanae* being predominant in post-weaned calves, while *C. andersoni* is the main infective species in adults [13, 16–22]. The latter three species have non or low pathogenic potential and seem to be host-adapted [23–25]. In contrast, *C. parvum* is the main source of infection in pre-weaned calves and the only species associated with typical cryptosporidiosis symptoms in cattle [7, 17, 19, 26, 27]. Most notably, *C. parvum* is capable of infecting multiple animal hosts and is the primary zoonotic agent of cryptosporidiosis [5, 28]. Hence, it is of particular concern that, during the infective stage, calves can shed an extraordinary number of oocysts in their feces (roughly 10 × 10^10^, per day) [29]. Since oocysts are extraordinarily resistant to a vast array of conditions, including common disinfectants, they can persist in the environment for long periods, and become readily infective after ingestion by humans and other animals [30–32].

To truly uncover the species diversity, zoonotic potential, and transmission dynamics of the different *Cryptosporidium* spp. circulating in cattle, molecular tools must be employed. By targeting and amplifying the *18S* ribosomal RNA gene followed by sequencing, it is possible to accurately distinguish and characterize *Cryptosporidium* at the species level [2, 33]. Subsequently, amplification of the 60 kDa glycoprotein gene (*gp60*) can be used to explore the intra-specific diversity of *C. parvum* in order to categorize it into different subtype families and further differentiate subtypes within the same family [2, 33, 34]. These molecular tools have been extensively employed to assess *C. parvum* role in zoonotic transmission and to trace sources of cryptosporidiosis outbreaks [6, 35]. *Cryptosporidium* spp. prevalence studies with further molecular characterization in cattle farms across Belgium, France and the Netherlands are quite sparse. Several reports have documented *Cryptosporidium* spp. occurrence and description of zoonotic species/subtypes in dairy or beef calves employing molecular tools in France [36–42]. To our knowledge, only one such study has been conducted in Belgium [43] and the Netherlands [44]. Therefore, the aim of the present study was to provide up-to-date information regarding the prevalence and transmission dynamics of *Cryptosporidium* species in cattle farms on a wide geographic area across Belgium, France, and the Netherlands, while subsequently investigating potential circulation of subtypes within and between farms and even countries.

## Material and Methods

### Geographical area of research and study model

This study was conducted under the Health for Dairy Cows (H4DC) project, funded by the Interreg 2 seas programme (https://h4dc-interreg2seas.eu/). This is a European territorial cooperation program covering the regions along the Southern North Sea and the Channel. This includes certain districts, such as the Flanders region of Belgium, the south of England, Les Hautes-de-France region in France, the west part of the Netherlands. The main objective of the project is to reduce the sanitary and economic impact of *Cryptosporidium* spp. on farms.

### Faecal sample collection

Faecal sample collection for all participating farms in this study occurred between September 2019 and June 2020. Fifty-seven farms were surveyed from three countries: Belgium, France and the Netherlands (**Figure 1**). The partners of the project selected the participating farms. Most farmers participated in this study because of one of the following reasons: (1) the dairy cows in their farms were encountering issues with diarrhoea; (2) they were willing to participate in the study regardless; and/or (3) they were willing to potentially change their farming practices. On each farm, veterinarians were instructed to collect 10 individual faecal samples, preferentially from the youngest calves up to three months of age and 10 individual faecal samples from their respective mothers, directly from the rectum and regardless of their clinical condition (diarrhoeic or non-diarrhoeic). Thus, on average, a total of 20 samples were collected per farm. For farms with less than 10 calves ≤ 3 months old, fewer samples of calves (and their mothers) were collected. All participating farms were visited once. In one of the farms in Belgium a total of 41 samples were collected (20 calves and 21 adults). Each sample was gathered using a single pair of disposable gloves, transferred to a 40 ml faecal collection container, and stored in a portable cooler until arriving to the storage place. Samples were stored at −20°C until finally shipped on ice packs to the laboratories for further analysis. All samples contained the animal identification number, the relation status between calf and mother, as well as, the date of sampling, age of the animal, farm identification and country of origin.

**Figure 1:**
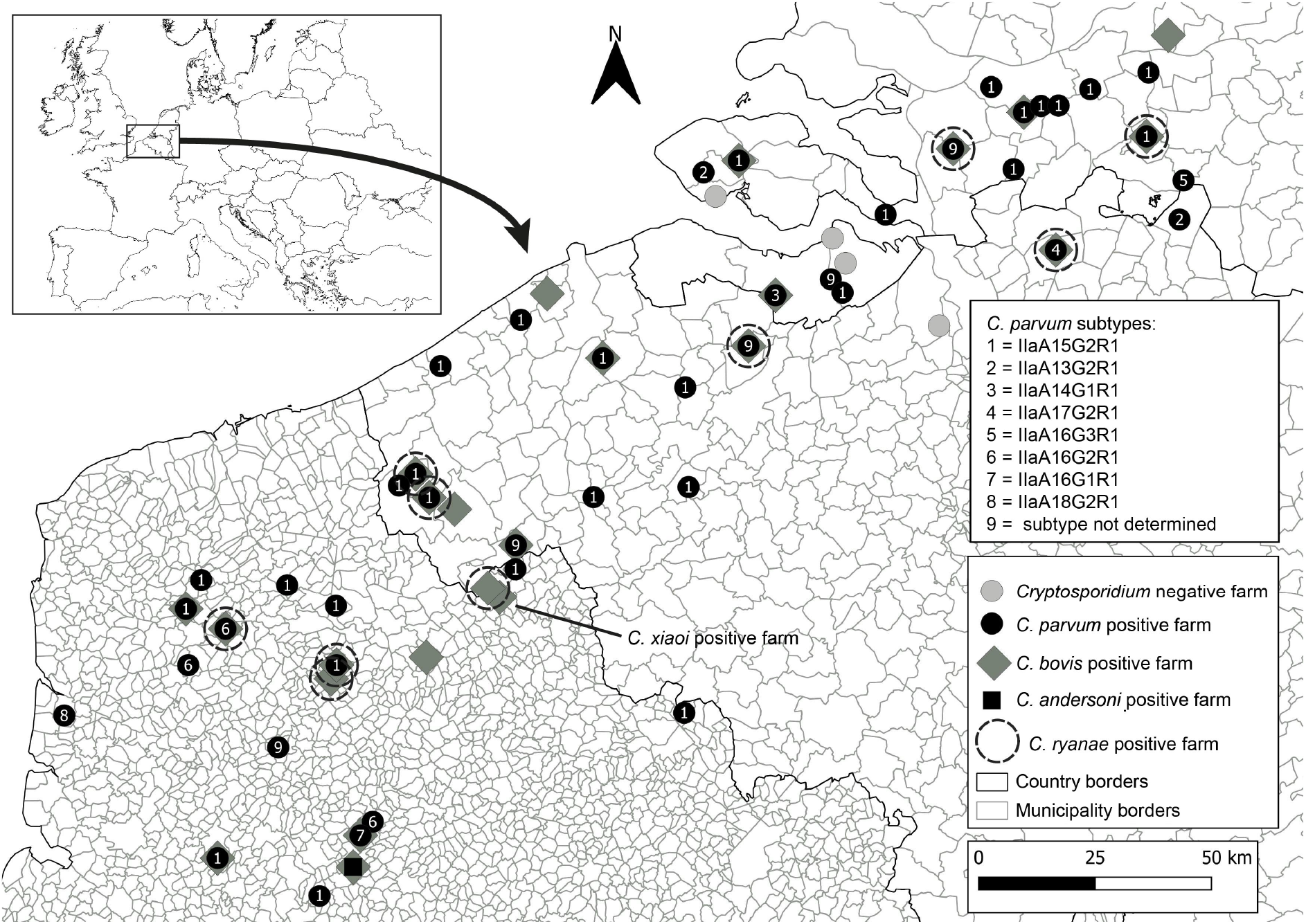
Geographical map of Belgium, France and the Netherlands indicating farms negative for *Cryptosporidium* spp. (grey circles) and farms positive for *C. parvum* (black circles), *C. bovis* (grey diamonds), and *C. andersoni* (black square). Distribution of *C. parvum* isolates with *gp60* subtypes are represented as numbers.

### Sample processing and DNA extraction

Frozen faecal samples were thawed overnight at 4°C and approximately 200 mg of faecal material was used to extract genomic DNA (gDNA) using the PureLink™ Microbiome DNA Purification Kit (Thermo Fisher Scientific, USA), according to the manufacturer’s instructions with slight modifications. After addition of clean-up buffer and instant vortex, samples were left to incubate at the fridge for 10 minutes to improve the removal of PCR inhibitors. DNA was then recovered with elution buffer and stored at −20°C until further analysis.

### *Cryptosporidium* spp. detection, and subtyping

*Cryptosporidium* spp. detection was attained through a nested PCR reaction amplification of the *18S* rDNA gene sequence. The primary reaction was conducted using the primers CRY_SSU_F1 5’-GATTAAGCCATGCATGTCTAA-3’ and CRY_SSU_R1 5’-TTCCATGCTGGAGTATTCAAG-3’ (product size: 723 bp) and the secondary reaction of the nested PCR was conducted using the forward primer CRY_SSU_F2 5’ -CAGTTATAGTTTACTTGATAATC-3’ and the reverse primer CRY_SSU_R2 5’-CCTGCTTTAAGCACTCTAATTTTC-3’ (product size ~631 bp) [45]. The primary PCR mixture was performed in a 25 μl volume containing 1 μl of template gDNA (concentration 10– 100 ng/μl), 0.4 μM of each primer, 12.5 μl of 2× PCRBIO Taq Mix Red (PCR Biosystems, UK). All amplifications were performed in a C1000 Touch PCR thermal cycler (Bio-Rad Laboratories, Inc., USA) with an initial denaturation at 94°C for 2 minutes, followed by 24 cycles of denaturation at 94°C for 50 seconds, annealing at 53°C for 50 seconds, and extension at 72°C for 1 minute. Lastly, a final extension step at 72°C for 10 minutes was also included. For the second PCR reaction, 1 μl of the PCR product obtained from the primary PCR reaction was used as template, and the remaining of the mixture was prepared as described above for the primary reaction. The cycling conditions for the second PCR reaction differed on the number of cycles, with 30 cycles instead of 24, and the annealing conditions, with 56°C for 30 seconds being used instead. Amplification of the 60 kDa glycoprotein gene (*gp60*) by nested PCR was carried out using the primers AL3531 5’-ATAGTCTCCGCTGTATTC-3’ and AL3535 5’-GGAAGGAACGATGTATCT-3’ (product size: 1000 bp) in the primary reaction of the nested PCR, and the primers AL3532 5’-TCCGCTGTATTCTCAGCC-3’ and AL3534 5’ - GCAGAGGAACCAGCATC-3’ (~850 bp) in the secondary reaction of the nested PCR [46]. Briefly, the primary and secondary PCR mixtures contained 2 μl of gDNA (ranging from 10 – 100 ng/μl) or of the primary PCR product, respectively, 0.2 μM of each primer, and 15 μl of 2× PCRBIO Taq Mix Red (PCR Biosystems, United Kingdom) in a total volume of 30 μl. The cycling conditions included an initial denaturation step at 94°C for 3 minutes, followed by 35 cycles of denaturation at 94°C for 45 seconds, annealing at 50°C for 45 seconds, and extension at 72°C for 1 minute. Lastly, a final extension step at 72°C for 7 minutes was also included. Nuclease-free water and gDNA extracted from 10^6^ of *Cryptosporidium parvum* IOWA strain purified oocysts (Waterborne™, Inc., New Orleans, LA, USA) were used as negative and positive control, respectively.

Secondary PCR products were separated by electrophoresis in a 2% (w/v) agarose gel (2%) stained with ethidium bromide (0.2 μg/mL), and visualised under a UV light system (Syngene G:BOX Chemi XX6, UK). Bands of interest were excised from the gel and the DNA was purified using Thermo Scientific GeneJET Gel Extraction Kit (Thermo Fisher Scientific, USA) following manufacturer’s instructions.

### Sequencing and phylogenetic analysis

Bi-directional sequencing of the purified secondary PCR products was outsourced to Eurofins (UK), who performed Sanger sequencing with the set of the primers used for the secondary PCR reaction. The quality of both forward and reverse nucleotide sequences generated for the expected amplicons were then manually assessed and trimmed if necessary, with ChromasPro version 2.1.9 (http://technelysium.com.au/wp/chromaspro/), assembled into a consensus sequence and mismatches corrected with the same software. The final assembled consensus sequence was then compared with GenBank reference sequences using the Basic Local Alignment Search Tool (BLAST) from the National Center for Biotechnology Information (NCBI) (http://blast.ncbi.nlm.nih.gov/Blast.cgi).

The sequences generated in this study were aligned with each other and with reference sequences from GenBank by MAFFT v.7 (https://mafft.cbrc.jp/alignment/server/) and the sequence alignment was manually inspected with BioEdit version 7.2.5 (https://bioedit.software.informer.com/). Phylogenetic analyses were performed and best DNA/Protein phylogeny models were selected using the MEGAX software [47, 48]. Phylogenetic trees were inferred using maximum likelihood (ML), with the substitution model that best fit the alignment selected using the Bayesian information criterion. The Tamura 3-parameter model [49], was selected for *SSU* and *gp60* alignments. Bootstrap support for branching was based on 1,000 replications. Phylograms were drawn using MEGAX and were manually adjusted using CorelDrawX7.

*Cryptosporidium parvum* allelic family and subtype was identified from the partial sequence of *gp60* gene based on the subtypes nomenclature reported previously [34]. The *18S* rDNA and *gp60* sequences derived in this study were deposited in GenBank. The accession numbers are: MW947282-MW947436 for *18S* rDNA of *C. parvum*, MZ021416 for *C. xiaoi*, MZ021417-MZ021429, MZ021431, MZ021433-MZ021436, MZ021438-MZ021441, MZ021443, MZ021445, MZ021447, MZ021449, MZ021451-MZ021461, MZ021464-MZ021467 and MZ021469-MZ021470 for *18S* rDNA for *C. bovis*, MZ021426 for *C. andersoni* and MZ021430, MZ021432, MZ021437, MZ021442, MZ021444, MZ021446, MZ021448, MZ021450, MZ021462-MZ021463, MZ021468 and MZ021471 for *C. ryanae*. The accession numbers MW996760-MW996892 are for the *gp60* of *C. parvum*.

## Results

### Sampling report

In this study, a total of 1,084 faecal samples from 57 farms were collected. These included 545 faecal samples from dairy calves and 539 from cows. Twenty of these were located in the Netherlands, 20 in France, and 17 in Belgium (**Figure 1**). For each farm, between 6 to 41 faecal samples were collected (median 20 and mean of 18.4 ± 4.4) and a roughly even ratio of faecal sample collection from calves and their corresponding mother cows was obtained. The age of the tested calves varied from 0 to 106 days (14 median and mean of 21.6 ± 21.1)

### *Cryptosporidium* spp. occurrence and prevalence in farms across Belgium, France and the Netherlands

Using amplification of a partial *18S* rDNA gene fragment with nested PCR revealed at least one animal tested positive for *Cryptosporidium* in 94.1% (n=17) farms in Belgium, 100% (n=xx) in France and 85.0% (n=17) in the Netherlands. Considering all farms, the overall occurrence of *Cryptosporidium* at farm level was 93.0% (53/57).

Within Belgium, *Cryptosporidium* spp. infections ranged from 5.0% (1/20 cows, farm 9) to 40.0% (8/20 cows, farm 11). Infection rates ranged between 5.6% (1/18 cows, farm 18) to 50.0% (3/6 cows, farm 31) in France, while in the Netherlands rates varied between 5.0% (1/20 cows, farm 51) to 30.0% (6/20 cows, farms 46, 50, 56 and 57) (**Table 1**).

**Table 1.**
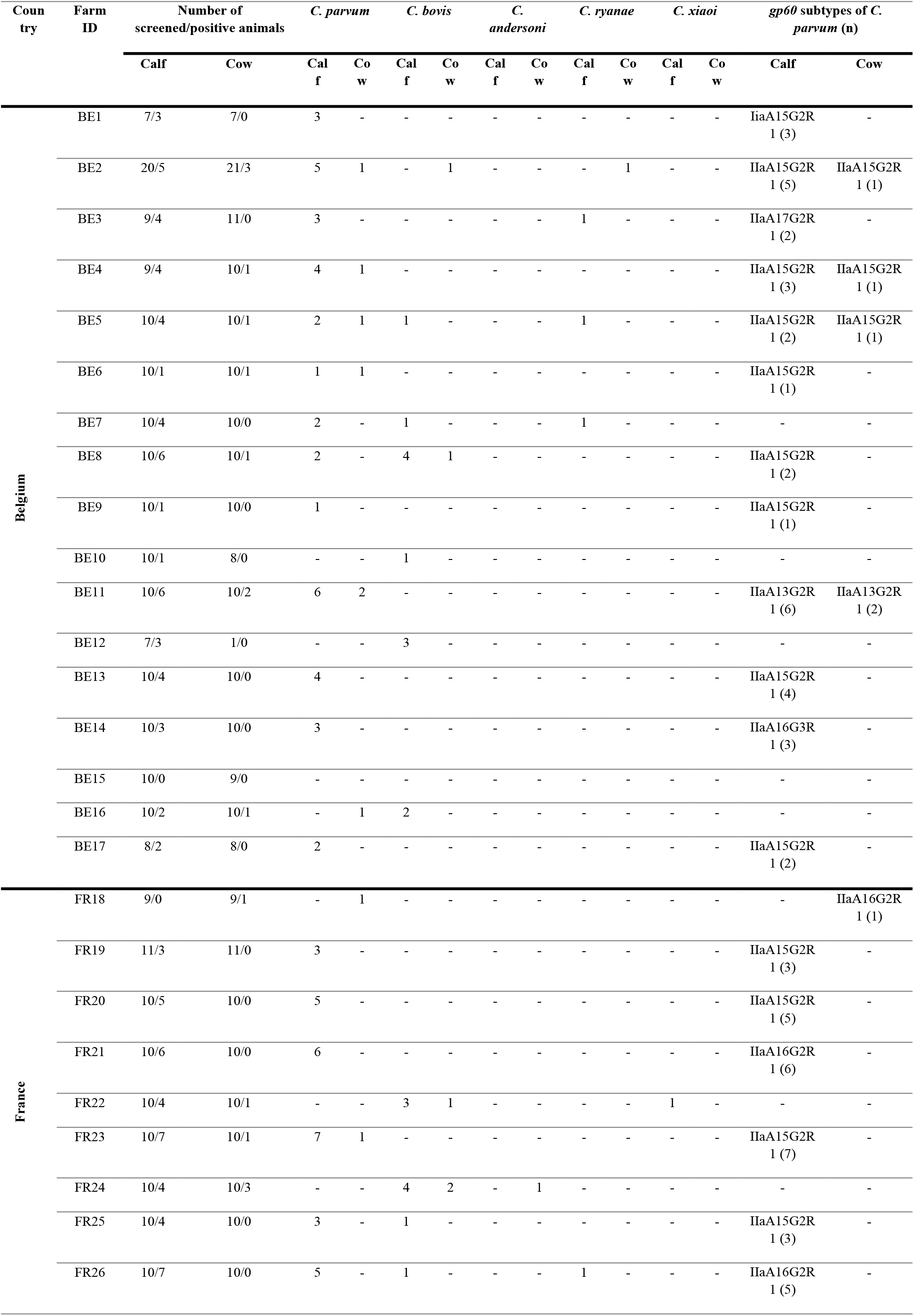
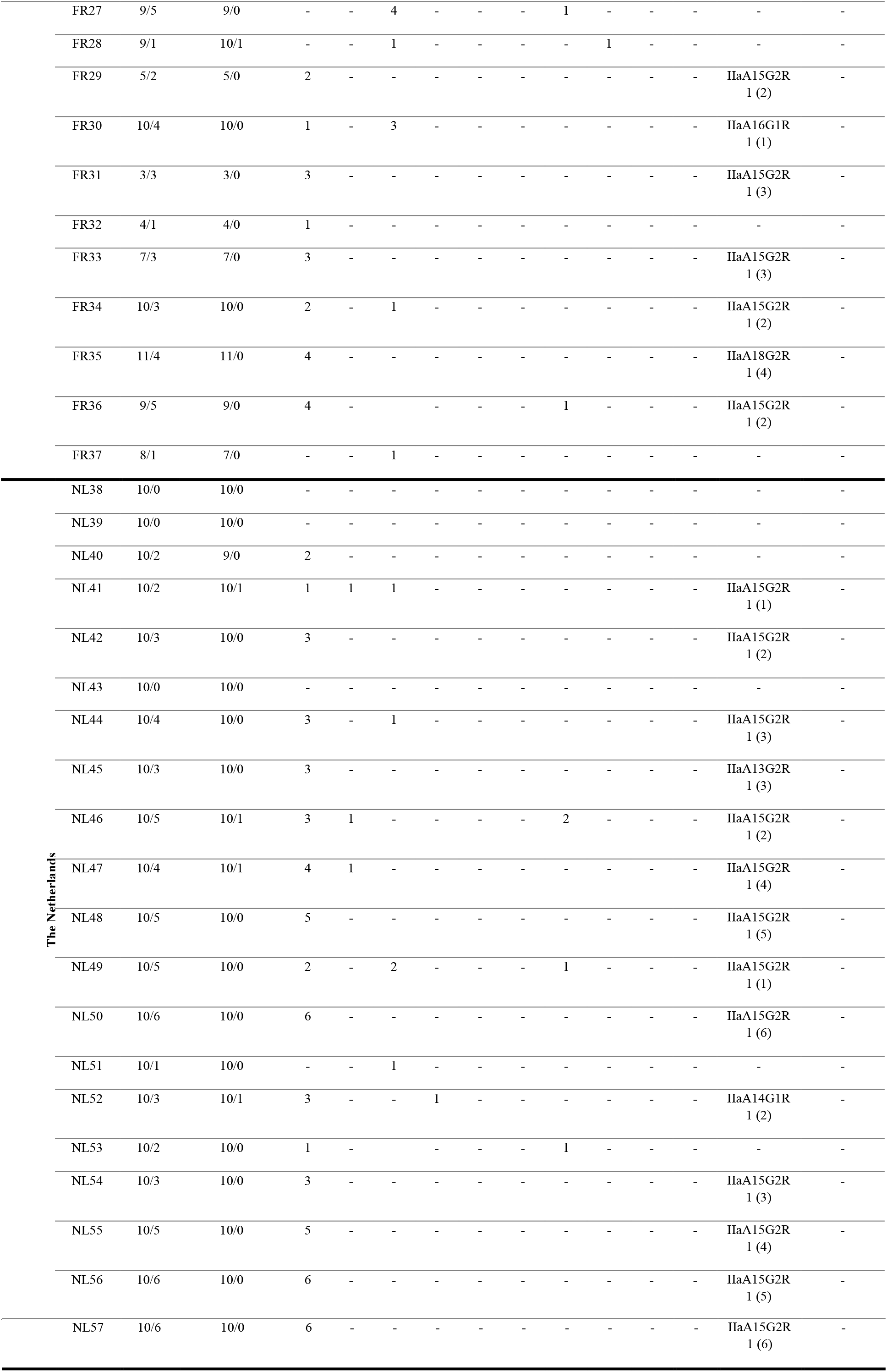
Distribution of *Cryptosporidum* spp. and *C. parvum* subtypes in all dairy cattle from the Netherlands, Belgium and France included in this study.

In terms of animals, 63 out of 335 were positive for *Cryptosporidium* spp. (18.8%) in Belgium, with young calves exhibiting a higher susceptibility (31.2%; n=170) to infection as opposed to adults (6.1%; n=165). In France, 79 out of the 350 (22.6%) animals tested positive with a prevalence of 41.1% (n=175) in calves as opposed to 4.0% prevalence in adults (n=175). Lastly, in the Netherlands’ farms, 69 out of 399 animals (17.3%) were positive, with 32.5% (n=200 prevalence in calves compared to adults, 2.0% (n=199) (**Table 2**).

**Table 2.**
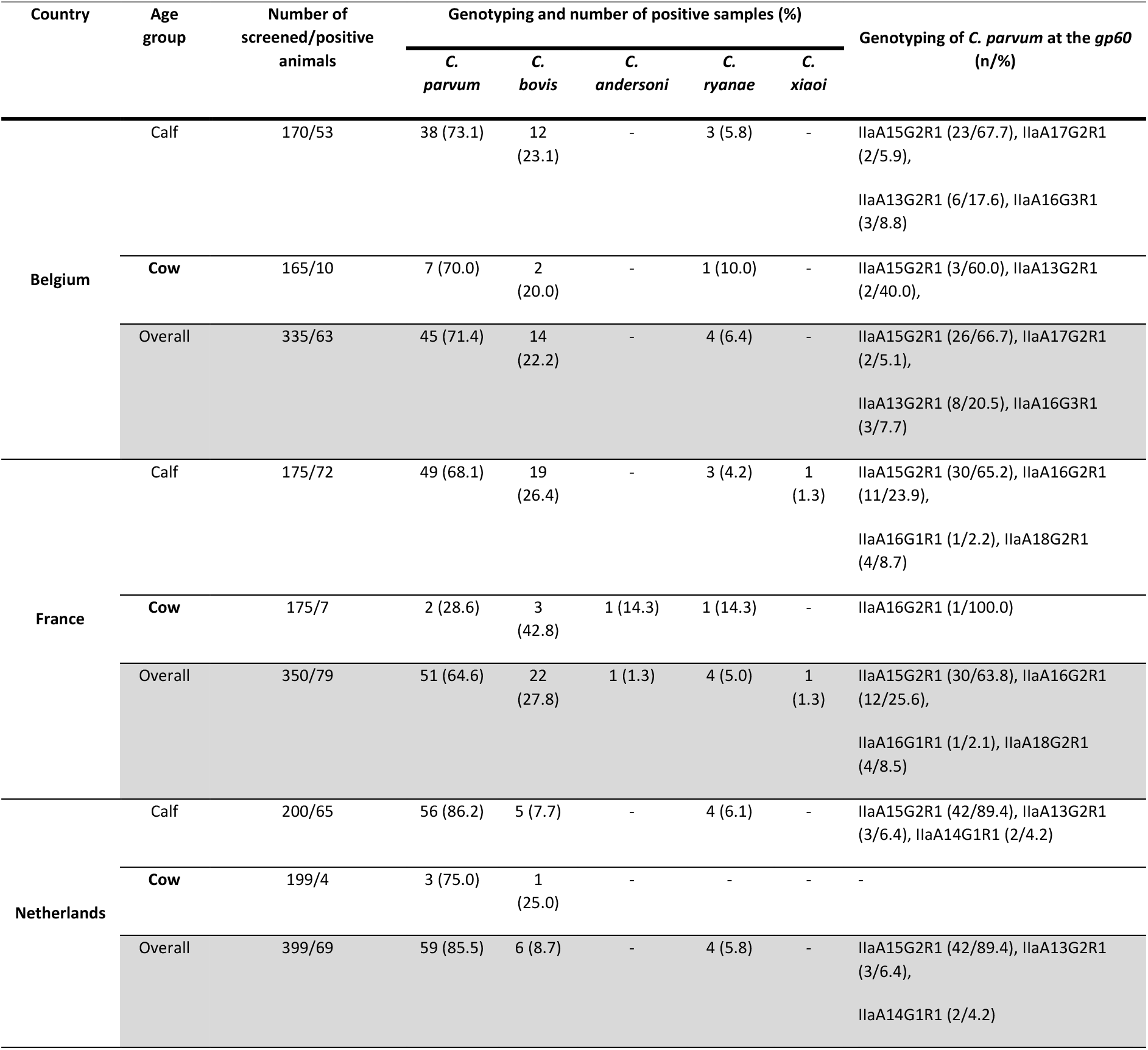
Prevalence and distribution of *Cryptosporidium* spp. and *C. parvum* subtypes in dairy cattle from the Netherlands, Belgium, and France.

#### Belgium

In Belgian farms, 63 animals were identified as *Cryptosporidium* spp. with 45 cases (71.4%) identified as *C. parvum*, 14 (22.2%) as *C. bovis* and four as *C. ryanae* (6.4%). In calves *C. parvum* was identified in 38 cases (73.1%), while 12 (23.1%) were shown to be *C. bovis*, and three (5.8%) were assigned as *C. ryanae*. In adult dairy cows, we identified seven positives (70.0%) for *C. parvum*, two (20.0%) for *C. bovis* and one for *C. ryanae* (10.0%) (**Table 2**). Sequence analysis of the *18S* rDNA for *C. parvum* isolates, revealed 100% nucleotide identity to the AH006572 reference sequence for 44 sequences (accession numbers MW947333-MW947340 and MW947342-MW947377) and 99% nucleotide identity to AH006572 reference sequence for the remaining sequences detected (accession number MW947341, displaying an A-to-T transversion at position 223, and an A-to-G transition at position 562). Sequence analysis of the *C. bovis* isolates, revealed 100% nucleotide identity to the AB777173 reference sequence for 12 sequences (MZ021445, MZ021447, MZ021449, MZ021451-MZ021457 and MZ021460-MZ021461) and 99% nucleotide identity to the AB777173 reference sequence for two sequences (MZ021458-MZ021459). *C. ryanae* isolates exhibited a 100% nucleotide identity to the reference sequence FJ463193 for four sequences (MZ021444, MZ021446, MZ021448 and MZ021450) **(Figure 2)**.

**Figure 2:**
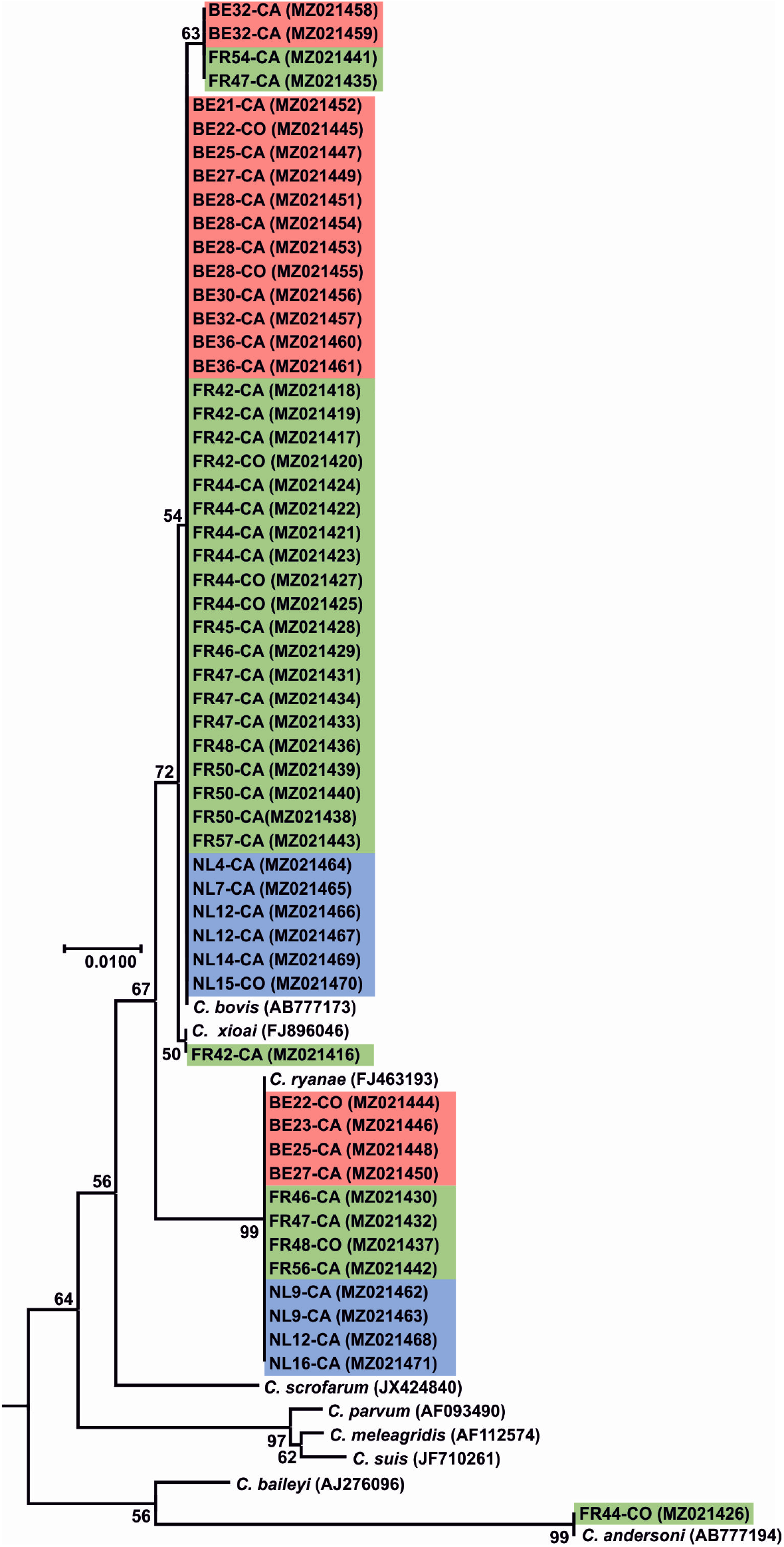
A maximum likelihood (ML) tree based on the *18S* rRNA gene sequences of *C. bovis, C. ryanae, C. xioai* and *C. andersoni* obtained in this study. Bootstrap values for the nodes with more than 50% support are shown. Sequences from this study are identified by country (NL for the Netherlands and highlighted in blue; BE for Belgium and highlighted in red; FR for France and highlighted in green), number of farm (e.g., NL4), and host age (CA for calf and CO for cow). The ML tree was rooted with an *18S* rDNA sequence from *Monocystis agilis* (accession number: AF457127). The GenBank accession number for each sequence is mentioned in parenthesis.

#### France

In France, a total of 79 samples tested positive for *Cryptosporidium* spp. with *C. parvum* being identified in 51 cases (64.6%), *C. bovis* in 22 (27.8%), *C. ryanae* in four (5.0%), *C. andersoni* in one (1.3%) and *C. xiaoi* in one (1.3%). In calves, 49 cases (68.1%) of *C. parvum* were detected, while *C. bovis* was identified in 19 cases (26.4%), *C. ryanae* in three (4.2%) and *C. xiaoi* in one (1.3%). In cows, *C. bovis* was observed in three cases (42.8%) *C. parvum* in two (28.6%), *C. andersoni* in one (14.3%) and *C. ryanae* also in only one case (14.3%) **(Table 2)**. Sequence analysis of *18S* rDNA for *C. parvum* isolates, revealed 100% nucleotide identity to the AH006572 reference sequence for 51 sequences (accession numbers MW947282-MW947332). Similar sequence analysis for *C. bovis* isolates, revealed 100% nucleotide identity to AB777173 reference sequence for 20 sequences (MZ021416-MZ021425, MZ021427-MZ021429, MZ021431, MZ021433-MZ021434, MZ021436, MZ021438-MZ021440 and MZ021443) and 99% nucleotide identity to the AB777173 reference sequence for two sequences (MZ021435 and MZ021441). *C. ryanae* isolates exhibited a 100% nucleotide identity to the FJ463193 reference sequence for four sequences (MZ021430, MZ021432, MZ021437 and MZ021442). Lastly, sequence analysis for *C. andersoni* isolate, revealed 100% nucleotide identity to the AB513856 reference sequence for the one sequence (MZ021426) while the *C. xiaoi* isolate (MZ021416) exhibited a 100% nucleotide identity to the FJ896046 reference sequence **(Figure 2)**.

#### The Netherlands

Out of the 69 samples shown to be *Cryptosporidium* spp. positive in the Netherlands, the majority of those (59 cases; 85.5%), were assigned as *C. parvum* while the remaining samples were identified as *C. bovis*, (6 cases; 8.7%) and *C. ryanae* (4 cases; 5.8%). In calves *C. parvum* was identified in 56 animals (86.2%), *C. bovis* in five (7.7%) and *C. ryanae* in four (6.1%). Out of the four positive cases in adult cows, three (75.0%) were identified as *C. parvum*, while the remaining one was *C. bovis* (25.0%) **(Table 2)**. Further sequence analysis of the *18S* rDNA gene for the *C. parvum* isolates, revealed 100% nucleotide identity to the AH006572 reference sequence for the 59 sequences detected in this study (accession numbers MW947378-MW947436). Sequence analysis for the *C. bovis* isolates, revealed 100% nucleotide identity to the AB777173 reference sequence for six samples (MZ021464-MZ021467 and MZ021469-MZ021470). Sequence analysis of the *C. ryanae* isolates presented a 100% nucleotide identity to the reference sequence FJ463193 for four cases (MZ021462-MZ021463, MZ021468 and MZ021471) **(Figure 2)**.

### *Cryptosporidium parvum* subtyping through *gp60* molecular analysis

Using *18S* rDNA sequencing and phylogenetic analyses, a total of 155 samples were identified as *C. parvum* positive. PCR products of the *gp60* gene were successfully obtained for 137 (88.4%) of these cases. Sequence analysis and subsequent subtyping revealed the presence of eight different subtypes belonging to the IIa subtype family **(Figure 3)**.

**Figure 3:**
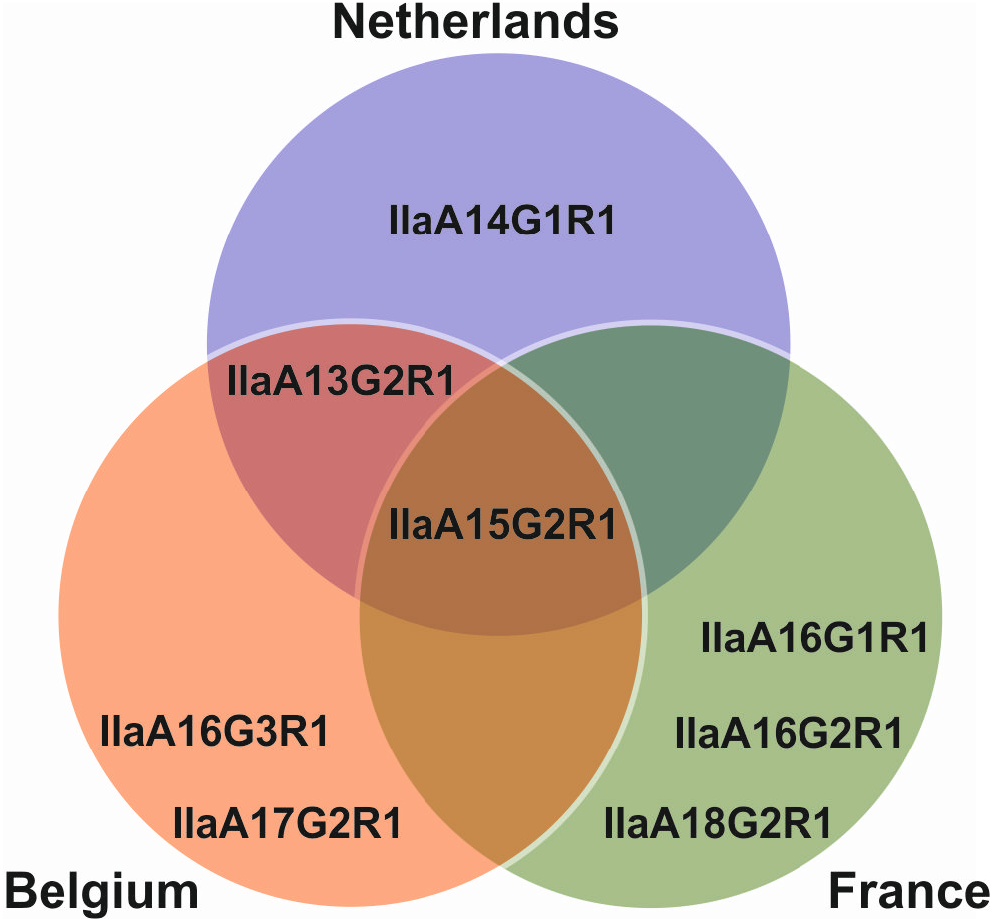
Venn diagram with all observed *C. parvum gp60* subtypes across Belgium, France, and the Netherlands.

At least one subtype of IIa family was found to circulate in 70% (n=20) of farms in the Netherlands, 70.6% (n=17) in Belgium and 70.0% (n=20) in France. Regarding, the distribution of subtypes between calves and cows, in the Netherlands all 47 (100%) of the subtypes identified were solely found in calves, while in Belgium 34 (87.2%) subtypes were described in calves and the remaining 5 subtypes (12.8%) in adults. Lastly, for French farms, 46 subtypes (97.9%) were identified in calves and only one subtype was identified in adults (2.1%) **(Table 1 and Table 2)**.

Subtype IIaA15G2R1 (100% identity to the reference sequence MK099855) was present in all three countries and was also the most widespread. In the Netherlands, Belgium, and France, this subtype was identified in 89.4% (42 out 47), 66.7% (26 out of 39), and 63.8% (30 out of 47) cases, respectively. In the Netherlands, this subtype was found in 70.6% (12 out of 17) farms which tested positive for *C. parvum*, while in Belgium and France this subtype was found in 64.3% (9 out of 14) and 60.0% (9 out of 15) farms, respectively. In the Netherlands, this subtype was only reported in calves, in 89.4% (42 out 47) infections, while in Belgium 67.7% (23 out of 34) calves and three out of five (60.0%) adults were found to have this subtype. Lastly, in France, this subtype was only detected in calves, in 65.2% (30 out of 46) infections.

The second most reported subtype in this study was IIaA16G2R1 (100% identity to the reference sequence MG516787), with all 12 isolates exclusively identified in France. This subtype was observed in 20% (3 out of 15) French farms that tested positive for *C. parvum*. Its presence was mainly in calves (23.9%; 11 out of 46 infections), with only one report in an adult cow. The third most abundant subtype was IIaA13G2R1 (100% identity with reference sequence, MN815775), with 11 isolates distributed through the Netherlands and Belgium. This subtype was found in just 1 out of the 17 (5.9%) Dutch farms which tested positive for *C. parvum*, and only in three calves out of 47 (6.4%). In Belgium, this subtype was also found in just one out of 12 (8.3%) farms which tested positive for *C. parvum*, with six out 34 (17.6%) calves and two out five (40.0%) cows testing positive for this subtype. The remaining five subtypes were: IIaA14G1R1, which was only observed in the Netherlands (99% identity to the reference sequence, AM937017 missing an adenine nucleotide at position 45); IIaA17G2R1 and IIaA16G3R1, which were only observed in Belgium (100% identity to the reference sequences, MG516783 and DQ192506, respectively); and IIaA16G1R1 and A18G2R1, which were only observed in France (100% identity to the reference sequences, KJ158747 and MK391451, respectively). All these subtypes were found in only a single farm and exclusively in calves (**Figure 1 & 4)**.

**Figure 4:**
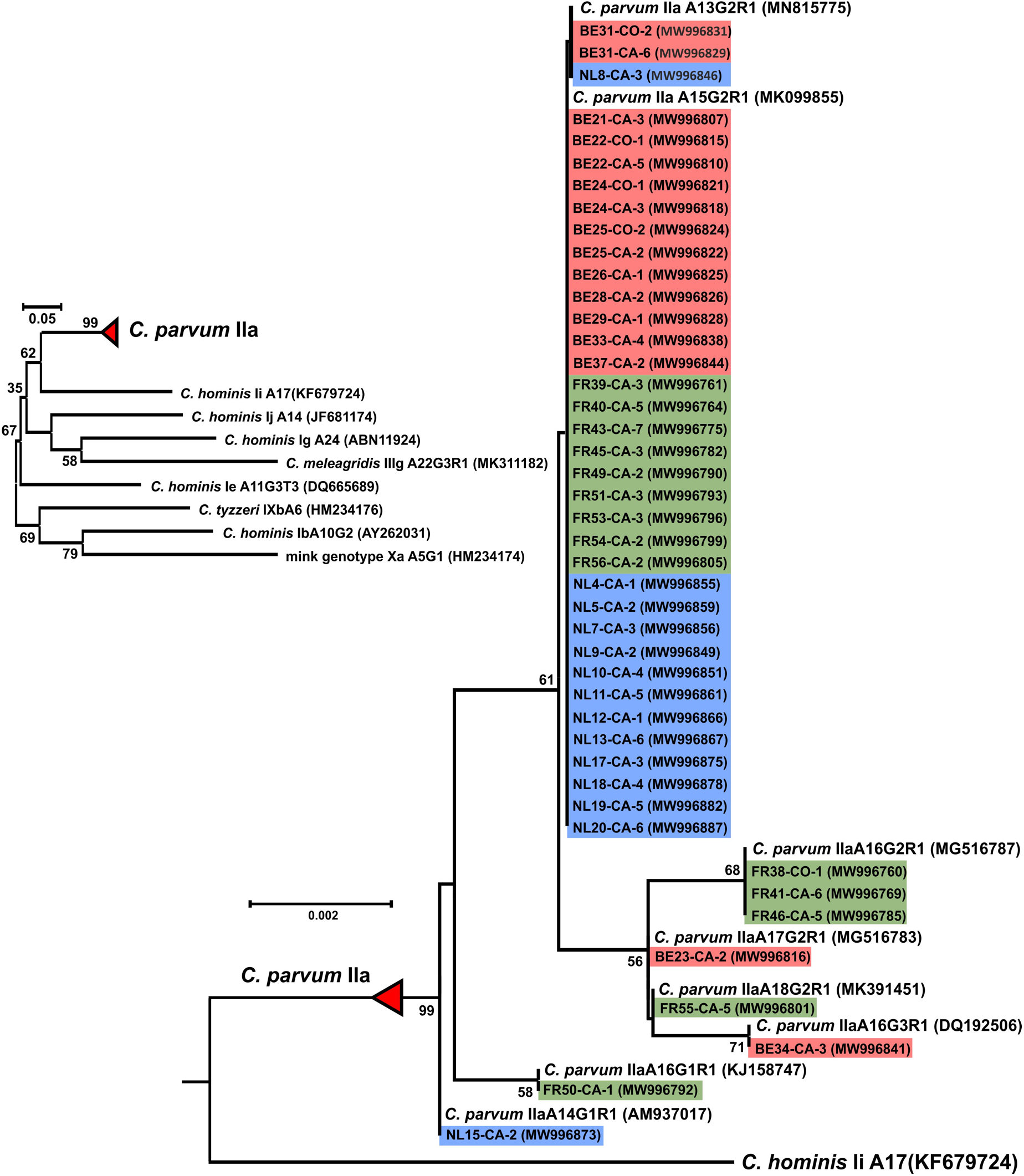
A maximum likelihood (ML) tree based on the *gp60* gene sequences. Bootstrap values for the nodes with more than 50% support are shown. *gp60* subtype family from this study is identified by country (NL for the Netherlands and highlighted in blue; BE for Belgium and highlighted in red; FR for France and highlighted in green), number of farm (e.g., NL4), host age (CA for calf and CO for cow), and the number of detected isolates of the subtype at the farm. The GenBank accession number for each representative sequence is mentioned in parenthesis.

## Discussion

*Cryptosporidium* is the causative agent of cryptosporidiosis in humans and other animals. In the farming industry, cryptosporidiosis is a major concern among livestock. *Cryptosporidium parvum*, in particular is considered to be a major cause of disease in neonatal calves, resulting in profuse diarrhoea and in extreme cases even death [13]. Herein, we employed molecular techniques based on *18S* and *gp60* gene analysis aimed to assess *Cryptosporidium* spp. prevalence as well as perform subtyping of *C. parvum* among cattle in several dairy farms distributed across three countries. Contribution of several factors in *C. parvum* spreading were also evaluated. Our analyses revealed that the average prevalence of *Cryptosporidium* spp. in screened farms, was 93.0%, with France having the highest at 100%, followed by Belgium at 94.1% and the Netherlands at 85.0%. These results are in line with previous epidemiological studies recorded in beef and dairy farms in France. For instance, Follet et al. (2011) reported presence of *Cryptosporidium* spp. in all examined beef farms following molecular detection using *18S* rDNA gene, while Mammeri et al. (2019) described presence of *Cryptosporidium* spp. on 92.3% of dairy farms using direct immunofluorescence assay (DFA) screening. Interestingly, in Belgium, the only published study in dairy farms reported presence of *Cryptosporidium* spp. on 32 out of 100 farms (32.0%) using quantitative immunofluorescence assay (IFA) for parasite detection [43]. In the Netherlands, the only study carried out to assess *Cryptosporidium* spp. prevalence in cattle did not provide any information on *Cryptosporidium* spp. per farm occurrence [44]. Similar prevalence studies in Europe, employing nested PCR targeting the *18S* gene, documented less overall occurrence of *Cryptosporidium* spp. with 44.5% in Spanish dairy and beef farms, 62.3% in dairy farms of Estonia [50, 51] and, in the Czech Republic, 79.2% in calves and 30.4% in cows [52, 53]. In Germany and Italy, following microscopy methods for *Cryptosporidium* spp. detection, it was also observed a lower prevalence of the parasite at farm level with 68.2% in Italian dairy farms and a 70.0% prevalence in German calf farms [54, 55]. A common issue between all these studies is the lack of consistent methods for sampling and parasite detection, which will allow accurate comparisons between them, while avoiding detections bias.

At individual level by country Belgium, France and the Netherlands totalled 18.8 %, 22.6 %, and 17.3 % prevalence respectively of *Cryptosporidium* spp. infection out of all sampled animals. Past studies in France reported higher *Cryptosporidium* spp. infections in beef and dairy calves (ranging from 41.5 to 100%) [36, 37, 39, 41, 42, 56]. Moreover, previous reports from Belgium also demonstrated slightly higher infections (37.0%) in dairy calves while, to our knowledge, no such information is available for the Netherlands [43]. In nearby European countries, such as Germany, United Kingdom, Austria, Spain, and Italy prevalence of *Cryptosporidium* spp. in beef and dairy calves assessed using molecular, immunological or microscopy screening, varied between 16.7% to 100% [51, 54, 55, 57, 58]. This data seems to correlate with equivalent findings worldwide where livestock parasite infections appear to lessen with the increase in age of cattle [19, 27, 59–65]. However, two recent studies carried out in dairy and beef farms in the United Kingdom pointed towards much higher infection rates in adult cattle with a reported *Cryptosporidium* spp. prevalence of roughly 80.0% in both studies, which was attributed to improved sensitivity in methods used during both investigative works [58, 66]. The observed variances in reported infections might stem from distinct geographical locations in association with climate variations but also due to differences linked to the study design, with factors such as sample size, age, herd size, total of farms investigated, farm management practices and screening methods applied playing an important factor [13].

Age-related infection predisposition to different *Cryptosporidium* species was also observed in our results after nested PCR analysis of the *18S* gene. *Cryptosporidium parvum* was clearly the dominant species in calves in all three countries, followed by *C. bovis*, *C. ryanae* and *C. xiaoi*, the latter having been observed in a single calf in France. Previous molecular studies conducted in dairy and beef calves in France and Belgium support our findings, with *C. parvum* being also the predominant infective species in pre-weaned beef and dairy calves [36, 37, 40, 43]. Other molecular studies across Europe also seem to indicate *C. parvum* as the dominant *Cryptosporidium* species in pre-weaned calves [50–52, 54, 58, 67]. The presence of *C. bovis* was less prominent in calves from this study, but this species has been reported in some parts of Europe and across the world as the major infecting species in beef and dairy calves [25, 68, 77, 69–76]. *Cryptosporidium ryanae* was sporadically detected in calves across the three countries, which seem to be consistent with other molecular studies performed in beef and dairy farms in France and in other parts of Europe [25, 36, 41, 50–52, 58, 74]. This species was, to our knowledge, observed for the first time in cattle from the Netherlands and Belgium. Both *C. bovis* and *C. ryanae* are mostly associated with infections in post-weaned calves and have yet been linked with cryptosporidiosis illness [13, 78]. *Cryptosporidium xiaoi,* the host-specific species for sheep [79], was only recorded in one calf in France herein. Reports of infection in cattle are scarce, however this species is reported occasionally in Jordan, China and Spain [51, 80, 81]. In Ireland infections in cattle were reported as *C. bovis/C. xiaoi* due to inability of distinguishing the two at the *18S* rDNA gene level [79, 82]. Previous cattle infections with *C. xiaoi* were either attributed to cross-infections between livestock sharing the same grazing fields or due to contact with contaminated water [51, 80]. To our knowledge, none of the farms included in this project had any association with other (small ruminants), thus the source of *C. xiaoi* remains unresolved.

The preponderance of *C. parvum* infections in calves has been debated before, with some studies suggesting that calves with *C. parvum* infections shed oocysts at a higher density when compared to other infective species, such as *C. bovis* or *C. ryanae*. The increased shed could lead to a preferential DNA amplification of the dominant *C. parvum* in a sample during *18S* rDNA gene-targeted PCR while, at the same time, masking the infection by other *Cryptosporidium* species and suppressing their detection [59, 83–85]. Previous molecular studies in cattle mention *C. andersoni* as the primary force behind *Cryptosporidium* infections in adult cows [18, 27, 59, 86]. However, in the present study, *C. parvum* was the dominant species infecting cows in the Netherlands (75%) and Belgium (70%), which was similar to what was reported in two recent studies carried out in United Kingdom cattle farms, which might indicate that adults shed *C. parvum* oocysts more frequently than previously thought [58, 66]. Additionally, reports from Spain also documented lower *C. andersoni* infections in adult cows in cattle farms and instead found *C. bovis* as the main species present in adult cattle, as observed in our study in France [51]. These results might indicate that *C. andersoni* is not as predominant in adult cattle as previously thought, and the established *Cryptosporidium* spp. age-related prevalence is not necessarily uniform across different regions.

To further explore the genetic diversity within *C. parvum* and assess potential zoonotic transmission, subtype assessment was undertaken through *gp60* locus analysis. In total, eight different subtypes were detected across the three countries and, notably, all belonged to the IIa zoonotic allelic family. The IIa allele is exceptionally prevalent in young cattle and has been linked with various occurrences of zoonotic transmission (both sporadic and outbreak cases) of *C. parvum* in Europe [6, 35, 87]. Subtype IIaA15G2R1 was found to be the most prevalent in all countries with the Netherlands reporting a total prevalence rate of 89.4%, Belgium at total of 66.7% and France a total of 63.8%. In western mainland Europe, the IIaA15G2R1 subtype was previously described as the most prevalent in cattle [36, 37, 40, 42–44]. Similar observations were reported in other European countries, including Portugal, Italy, Spain, Germany, Czech Republic, Austria, Slovenia, and the United Kingdom [20, 51, 52, 54, 55, 57, 66, 67, 88] and in other parts of the world such as Japan and Uruguay [89, 90]. IIaA15G2R1, is deemed as the most prevalent IIa subtype in humans in developed countries, with several cases of human cryptosporidiosis linked to it [6, 44, 67, 87, 88, 91–93]. Several previous outbreaks with this subtype in the United Kingdom were traced back to contact between farm animals and humans pointing towards zoonotic potential [35]. In fact, it has been speculated that this subtype is hyper-transmissible, which might explain its preponderance in zoonotic infections across the world [3].

The second most documented subtype was IIaA16G2R1, with all occurrences being detected in French farms, and with a prevalence of 25.6%. This subtype was previously described in cattle of France with a much lower prevalence of just 3.9% [36]. Interestingly, this subtype was also previously observed in cattle located in Belgium and the Netherlands [43, 44]. Cattle infected with this subtype was also previously found in other parts of Europe, including Portugal, Germany, Spain and Estonia [50, 55, 88, 94]. IIaA16G2R1, has been known to cause sporadic human cryptosporidiosis in a few countries, namely New Zealand, Canada, Spain, and Jordan [91, 92, 95–97].

The third most abundant subtype in our study was IIaA13G2R1, which occurred only in the Netherlands and Belgium, with a total prevalence of 6.4% and 20.5%, respectively. This subtype has been found previously in both countries at lower frequencies than the ones reported herein, with a prevalence of 1.5% in both the Netherlands and Belgium [43, 44]. From all other European countries, so far this subtype has only been reported in the United Kingdom [20]. This *C. parvum* subtype appears to be uncommon among calves with only Algeria, Turkey and Canada reporting its occurrence [98–100]. IIaA13G2R1 also has a low prevalence in humans with confirmed cases in Malaysia, South Korea, New Zealand, and Canada and thus its zoonotic potential is considered low [91–93, 101].

The remaining five subtypes detected in this study had a low prevalence in all the studied countries, with less than five occurrences in each. To our knowledge, subtype IIaA14G1R1, which was only detected in the Netherlands, was for the first time described in this country and its lower prevalence (4.2%) seems consistent to other reported frequencies in cattle across various European countries such as Germany, Poland, Austria, and Estonia [50, 55, 57, 102]. Cryptosporidiosis cases in humans for this subtype have been documented in New Zealand, Canada, Slovenia and Slovakia [67, 91, 92, 103]

Subtypes IIaA16G3R1 and IIaA17G2R1, were only found in Belgium and were for the first time described in this country, with a 7.7% and 5.1% prevalence, respectively. Subtype IIaA16G3R1 presence was previously described in France and the Netherlands, with similar frequencies [36, 40, 44]. Subtype IIaA17G2R1 was only observed before in the Netherlands with also with a similar frequency to that observed in this study, though two studies conducted in France river and sea waters reported the presence of this subtype on fish [44, 104, 105]. Subtype IIaA16G3R1 has been found in cattle in several European countries, such as Germany, Italy, Spain, Poland, and the United Kingdom [20, 51, 54, 55, 102, 106]. Subtype IIaA17G2R1 has been found in cattle in Slovakia, Poland, Spain, Germany, and the United Kingdom [20, 55, 94, 102, 106, 107]. Both subtypes have been previously observed in humans in New Zealand, Canada, Denmark, and Iran [87, 91, 92, 97, 108]

Lastly, subtypes IIaA18G2R1 and IIaA16G1R1 were found only in France with a prevalence of 8.5% and 2.1%, respectively. Similar studies in France found both subtypes present in calves with one study reporting a similar prevalence of IIaA16G1R1 to the one reported in this present study [36], while the another study found the IIaA18G2R1 to be the most prevalent in beef calves [41]. Subtype IIaA16G1R1 has also been observed before in the Netherlands, with a similar prevalence to the one reported in our study [44]. Subtype IIaA16G1R1 has also been reported in cattle from other European countries, including Sweden, Czech Republic, Hungary, Poland, Romania, Slovenia, Germany and Estonia, [50, 52, 55, 67, 69, 74, 75, 102, 109, 110], while subtype IIaA18G2R1 has been described before in cattle from Germany and the United Kingdom [58, 66, 106, 111]. Human infections with these subtypes have also previously been detected in New Zealand, Canada, Sweden, Slovakia, Slovenia, Estonia, and the United Kingdom [35, 67, 91, 92, 103, 112, 113].

Through this study, only one subtype per farm was observed which might indicate endemicity at the farm level confirming results from previous studies [40, 67, 102]. This lack of genetic diversity may not be the rule though, with recent reports finding that more than one subtype per farm might be the norm [20, 37, 50, 54, 55]. The lack of genetic diversity per farm in our study might stem from the approach used, since conventional *C. parvum gp60* subtyping with Sanger sequence only targets the dominant subtype. The application of next-generation sequencing (NGS) and/or single cell genomics [114–116] could provide a more reliable way to unearth the real multiplicity of *C. parvum* within a herd and within the same host. In fact, two reports using NGS described up to ten individual subtypes per sample in cattle and human isolates [117, 118].

Another aim of this study was to investigate the possible role of mothers in transmitting the parasite to their new-born calves. Previous investigations on the subject were conducted before the advent of molecular subtyping tools thus no conclusions could be made. A recent paper addressed this issue with *gp60* molecular subtyping [58]. The subtype analysis in our study and in agreement with previous *18S* rDNA data analysis, yielded a much higher amount of *C. parvum* genetic information in calves than in adults. After *gp60* analysis, only two pairs of adults-calves were found to share the same *C. parvum* subtype, one pair in farm 5 and the other in farm 11, both located in Belgium. This suggests that there is no clear link for maternal transmission of the parasite. A similar and recently published work, also looked into this hypothesis and did not find a strong link in the spread of *C. parvum* between adults and calves, finding that calves and adults shed different *C. parvum* subtypes [58]. However, it is possible that standard *gp60* molecular and Sanger sequencing methods employed in this study might not have been discriminatory enough to uncover the true diversity of subtypes due to bias [116–118]. Moreover, sample collection occurred at a single time point instead of several. Thus the possibility of various additional subtypes going undetected due to differential shedding cycles cannot be excluded [58]. Although no strong evidence could be gathered regarding the source of infection of *Cryptosporidium* spp. within the farms, recent studies have started to investigate alternatives sources of transmission from outside the farms. For instance, zoonotic *C. parvum* subtypes sampled in cattle from UK farms were identical to those circulating in wildlife nearby, particularly in birds and deer [20, 66]. Remarkably, the same zoonotic *C. parvum* was also found in water bodies within/nearby the farms thus water might be an alternative route of *Cryptosporidium* transmission in cattle and, eventually, transmission to humans. Thus farmers should consider implementing better water and farm management systems to prevent further environmental spread and contamination [20, 66]. Moreover, although *C. parvum* transmission and infection is largely associated with mammals, recent studies conducted in aquatic environments highlighted its presence in edible fish (sea and freshwater), raising awareness for the importance of cattle as a source of environmental contamination and dispersion of *Cryptosporidium* spp. oocysts, since most of the fish were affected by the *C. parvum* subtype IIa [104, 105]. Therefore, suitable farm practises that improve animal well-being and decrease occupational risks to humans in close contact with cattle should be implemented while better farm management systems to prevent further environmental spread and contamination should also be applied.

## Conclusions

This study provides a detailed view of *Cryptosporidium* spp. prevalence across dairy cattle farms in Western Europe and is the first study in dairy farms in the Netherlands. Our findings indicate that *Cryptosporidium* spp. is widespread across dairy farms, with zoonotic *C. parvum* being the dominant species detected in calves across all the three countries included in this study. In addition, all the *C. parvum* subtypes identified in this study have been linked with cryptosporidiosis in humans, highlighting the potential of cattle as a reservoir for *C. parvum*. Our study also provides evidence that it is unlikely that adult cattle, plays a role as a source of infection, as sharing of *C. parvum* subtypes between adults and calves were only documented in two cases. However, it is important to note that other sources of infection could not be ruled out. Thus, follow-up studies should be conducted to assess a cross-borders and worldwide *Cryptosporidium* spp. prevalence, epidemiology, and transmission while considering a one-health approach on tackling cryptosporidiosis.

## Acknowledgments

This project has received funding from the Interreg 2 Seas programme 2014-2020 co-funded by the European Regional Development Fund under subsidy contract No 2S05-043. We would like to thank the farmers in the 2-seas region for volunteering and working with us on this project. Lastly, we would like to thank Dr. Eleni Gentekaki from Mae Fah Luang University (Thailand) for providing us constrictive feedback on the manuscript.

